# An Open-Source, Programmable Pneumatic Setup for Operation and Automated Control of Single- and Multi-Layer Microfluidic Devices

**DOI:** 10.1101/173468

**Authors:** Kara Brower, Robert Puccinelli, Craig J. Markin, Tyler C. Shimko, Scott A. Longwell, Bianca Cruz, Rafael Gomez-Sjoberg, Polly M. Fordyce

## Abstract

Microfluidic technologies have been used across diverse disciplines (*e.g.* high-throughput biological measurement, fluid physics, laboratory fluid manipulation) but widespread adoption has been limited due to the lack of openly disseminated resources that enable non-specialist labs to make and operate their own devices. Here, we report the open-source build of a pneumatic setup capable of operating both single and multilayer (Quake-style) microfluidic devices with programmable scripting automation. This setup can operate both simple and complex devices with 48 device valve control inputs and 18 sample inputs, with modular design for easy expansion, at a fraction of the cost of similar commercial solutions. We present a detailed step-by-step guide to building the pneumatic instrumentation, as well as instructions for custom device operation using our software, Geppetto, through an easy-to-use GUI for live on-chip valve actuation and a scripting system for experiment automation. We show robust valve actuation with near real-time software feedback and demonstrate use of the setup for high-throughput biochemical measurements on-chip. This open-source setup will enable specialists and novices alike to run microfluidic devices easily in their own laboratories.

Specifications table

**Specifications table**
|  |  |
| --- | --- |
| Hardware name | An Open-Source, Programmable Pneumatic Setup for Operation and Automated Control of Single- and Multi-Layer Microfluidic Devices<br>Geppetto (Python-based solenoid valve control software with GUI)<br>Geppetto CLI (Python-based solenoid valve software with command line scripting for experiment automation) |
| Subject area | • Engineering and Material Science<br>• Biological Sciences (*e.g.* Microbiology and Biochemistry)<br>• General |
| Hardware type | Biological sample handling and preparation |
| Open Source License | Pneumatics Instrumentation: GPL License<br>Geppetto Software: MIT License |
| Cost of Hardware | Variable, dependent on user customization<br>Full Build: \$10,380.<br>Abridged Build: \$4,455.<br>Manual Build: \$2,110.<br>Additional Modification Builds Costs shown in Table 1 |
| Source File Repository | Github: github.com/FordyceLab/Open_Source_Pneumatics<br>Supplementary Materials (In Article) |

## 1. Introduction

Microfluidic technologies enable the precise control and manipulation of fluid at the sub-millimeter scale [1]. Simple, single layer microfluidic devices with channels etched in glass or patterned into soft polymers (*e.g.* polydimethylsiloxane, or PDMS) have facilitated the study of small-scale fluid phenomena, automated sample loading and common laboratory tasks, and been used for high-throughput compartmentalization of biological reactions [2-9]. More complex multilayer microfluidic devices with monolithic soft polymer valves (also known as Quake-style valves [10,11]) have empowered additional research and commercial applications, including on-chip cell culture and tissue engineering [12], crystallization condition scanning [13-15], single cell analysis [16-18], high-throughput biochemical analysis [19-21], hydrogel engineering [22-24], and complex fluid handling tasks [11,25].

These multilayer microfluidic devices are typically fabricated in a two-layer geometry, where each layer is made by conventional soft lithography [10]. The ‘flow layer’ contains channels to load and transport samples. The ‘control layer’ contains a series of channels that create and control pneumatically-actuated valves. The two layers are separated by a thin polymer membrane. A valve is formed when flow and control channels of an appropriate size and shape cross [26]. When a control channel is pressurized, the thin membrane deflects into the crossing flow channel to modulate its flow. Both layers require their own set of input lines to interface with on-chip inlets. For complex chips with many control valves, a computer-controlled bank of solenoid valves is typically used to automate control of applied pressure to the valves.

While multilayer devices can serve as powerful tools for a broad range of biological systems [5,27-29], their widespread adoption has been limited, likely because few detailed open-source protocols and designs for building an automated pneumatic control systems have been published [3,30,31].

Previously, our group worked in conjunction with the Stanford Microfluidic Foundry to publish an open protocol that describes in detail how to fabricate custom multilayer microfluidic devices [32]. In this report, we continue the project of disseminating how-to knowledge on microfluidics by offering designs and detailed step-by-step build instructions for a flexible pressure-driven pneumatic setup that can control both fluid flow and integrated on-chip valves.

The primary pneumatics control system described here allows for automated, programmable control of single and multi-layer microfluidic devices with 48 control lines and 18 individually-addressable flow lines. We present customization options and instructions to build setups from 8 – 96 control lines and 6 – 48 flow lines using this base build. This setup confers several advantages to commercial alternatives including:

- flexible application programming and automation
- remote control to allow experiments to be controlled and monitored outside the lab
- extremely fast valve response times (using Modbus-pulsed solenoid valves)
- very low cost-per-input ratios (both control and flow channel inputs)

The build is modular and easy to customize for a user’s specific needs. We present two modified builds in a later section of the paper for lower complexity control, including a low-cost fully manual control configuration. Previous versions and variants of this setup, originating from the early days of the Stanford Microfluidic Foundry [33], have been used in tens of laboratories resulting in over a hundred peer-reviewed publications ([12,20-23,34,35] are some examples). However, full build details have never been published. Here, we describe the latest optimization of the pneumatic control system in detail to provide a complete tool kit that will allow even labs with little engineering expertise to build all of the instrumentation required for automated microfluidic control.

## 2. Hardware description

The open-source pneumatic setup and software (**Figure 1**) described here can be used for both manual and fully programmable pneumatic control of most PDMS-based microfluidic devices. Our associated software, Geppetto, has both an interactive GUI for manipulating device valves and visualizing their current state (open or closed) as well as command line scripting functionality for automating experiments.

**Figure 1.**
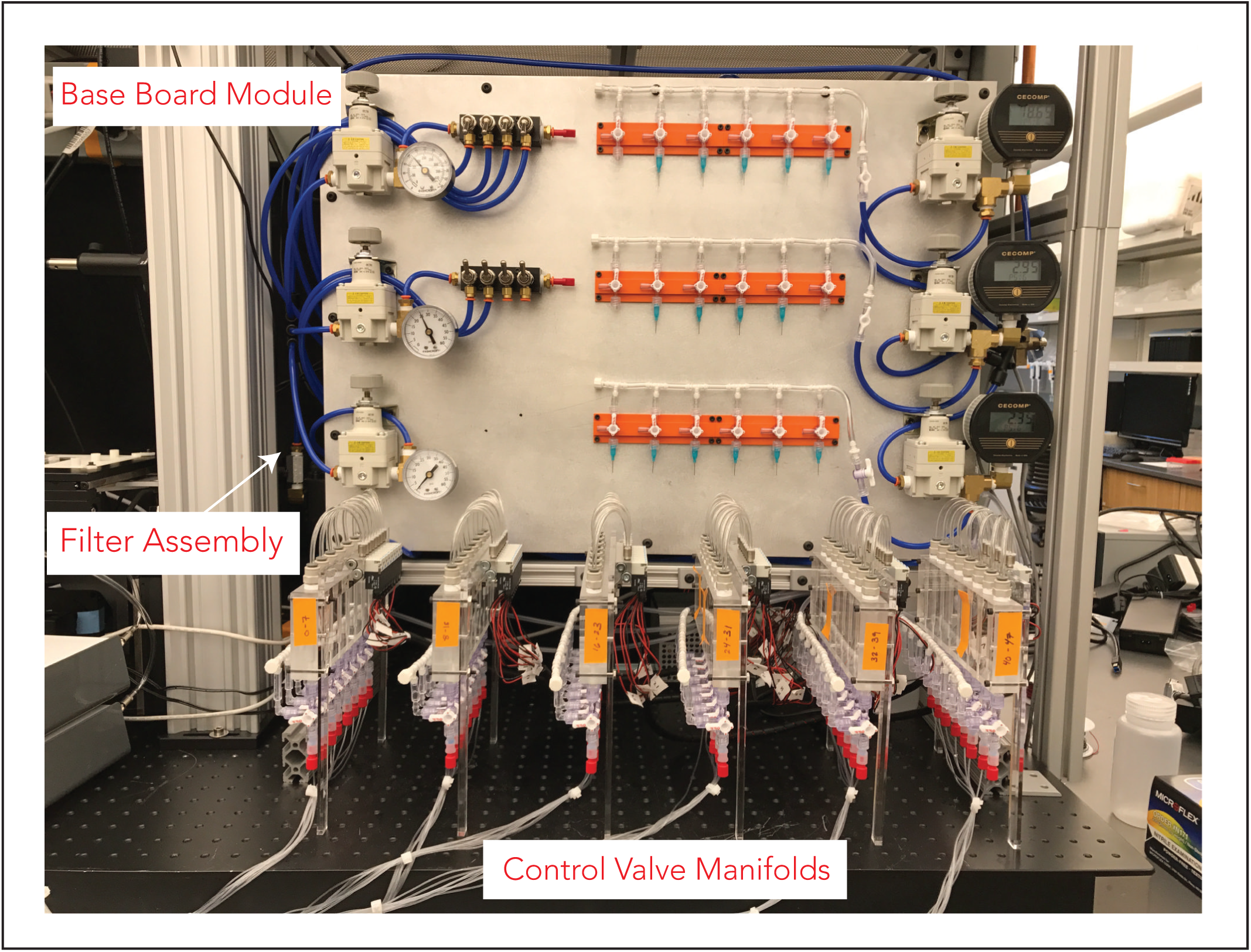
Photograph of a completed pneumatic build (Option 1) with all 5 build modules.

We present 3 build options for the microfluidic pneumatic control system, as listed below. This setup can additionally be expanded for up to 96 control lines and up to 48 flow lines. Control inputs need not correspond to the actual number of valves on the device - only the number of valves needing independent actuation. Typical multilayer microfluidic devices operate with 10 – 15 control lines and 10 – 15 sample lines, even for complicated tasks, but this build can accommodate devices with more or fewer inputs or the need to run multiple devices simultaneously on the same setup.

### 2.1. Build Options

In this paper, we present 3 options for building a microfluidic pneumatic setup:

**Option 1:** A full build for 18 flow lines and 48 individually addressable solenoid valves to drive 48 control lines for fully automated, programmable and remote operation of complex devices or multiple devices
**Option 2:** An abridged build for 6 flow lines and 8 individually addressable solenoid valves to drive 8 control lines for fully automated, programmable and remote operation of simpler devices
**Option 3:** A low-cost build for manual control of 18 flow lines and 12 control lines

The main text will describe the build for **Option 1**. The build is modular and can be easily modified to the simpler builds of **Options 2** and **3** to reduce time and cost, if only limited or manual control of microfluidic devices is needed.

### 2.2 Module Overview

The pneumatics control system is built in 5 **Modules** which are divided by manual and automated control capabilities. **Options 1** and **2** require all 5 **Modules**. The manual build, **Option 3**, requires a modified version of only **Modules 1** and **2**. The builds for **Options 2** and **3** are presented in a **Modifications and Customization Section.**

The Modules are as follows:

- **Module 1:** Base Board, Pressure Regulator Manifolds, and Flow Lines
- **Module 2:** Filter Assembly
- **Module 3:** Control Solenoid Valve Manifolds
- **Module 4:** Control Reservoir Loading Assembly
- **Module 5:** Digital Modbus Control

#### Manual Control Modules

This pneumatic setup employs pressure-driven flow control. Microfluidic devices, in general, can be operated either by pressure-drive flow (using constant pressure at the input) or volumetric flow (forcing a set flow rate at the input, as a syringe pump does). Volumetric flow results in unknown pressures inside the flow channels, which makes it difficult to ensure that on-chip valves close reliably and can lead to device delamination. This setup uses pressure-driven flow because it requires little additional equipment beyond common laboratory compressed air sources and provides robust valve actuation on-chip.

Controlling pressure-driven flow rates and actuating on-chip pneumatic valves requires a device-specific pressure range: too little pressure and valves don’t close properly, too much pressure and devices can delaminate or become otherwise damaged. Compressed air (from a house air supply, cylinder or compressor) provides a convenient and quiet pressure source but must be adjusted to the correct pressure for microfluidic devices using regulators. **Module 1** (Base Board, Pressure Regulator Manifolds, and Flow Lines) describes the base build for the pneumatic assembly to control input pressures. It contains a base board that mounts regulators for controlling both flow lines and control line manifold pressures as well as fluidic components for sample introduction or manual control of integrated on-chip valves.

**Module 2** describes how to connect a filtering assembly to the compressed air source to prevent device contamination and/or clogging of the air regulators.

#### Automated Control Modules

Automation of microfluidic valve actuation enables a wide range of additional assays and remote device control but requires additional control hardware. To control on-chip valves, we employ 24V, three-way, normally open solenoid valves which toggle between an ‘open’ pressurized state (with pressure from the control manifold regulators in **Module 1**) and a ‘closed’ atmospheric pressure state to close or open on-chip valves, respectively. These solenoid valves are used to pressurize or depressurize water within a reservoir directly connected to control lines, thereby providing individually-addressable control of each control line. Groups of eight of these solenoid valves are coupled to a plastic manifold containing eight water reservoirs connected to eight chip control lines. We call this assembly a control manifold; these manifolds are built in **Module 3** (Control Solenoid Valve Manifolds). **Module 4** (Control Reservoir Loading Assembly) adds on a large volume water bottle for easy loading and refilling of the water reservoirs in the control manifold. Finally, **Module 5** (Digital Modbus Control) describes the electrical connections between the solenoid valves and a microcontroller (the ‘WAGO’ fieldbus controller) to allow values to be digitally controlled by the user via a computer.

#### Pneumatic Control System Operation

Combined, these modules constitute all the components and connections necessary to provide pressure-driven flow control and actuate integrated on-chip valves of microfluidic devices. Initial setup of the system for experimental use is described in the **Operation Device Guide**.

**Figure 1**. Photograph of a completed pneumatic build (Option 1) with all 5 build modules.

**Figure 2**. Overall setup schematic with parts designated by letter and number (as referenced to the Bill of Materials).

**Figure 2.**
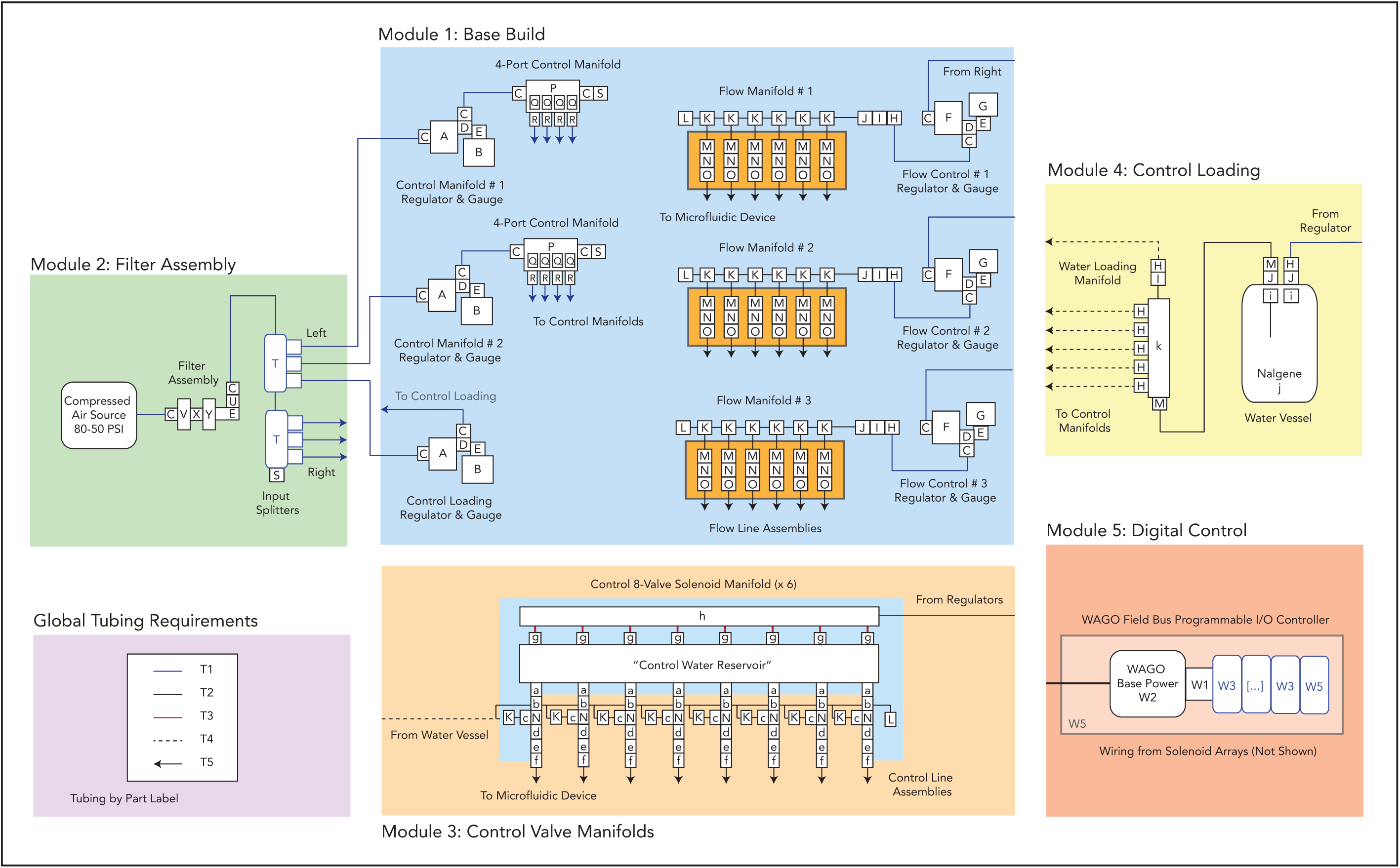
Overall setup schematic with parts designated by letter and number (as referenced to the Bill of Materials).

While at first glance this setup may seem complicated, we have built multiple setups with researchers of all technical backgrounds with little to no engineering training. Each module is accompanied by a schematic, parts list and illustrated step-by-step build instructions (**Build Guides**) that are included in the **Supplemental Material** of this report. The total build takes 1.5 days from start to device testing and is easily managed with 1-2 researchers and common laboratory tools.

### 2.3. Advantages of an Open-Source Build

Building an open-source pneumatic microfluidic control system has three main advantages to both academic researcher and small companies. First, open source builds are flexible and can be easily modified for a laboratory’s needs [30,36]. For instance, this paper describes two easily modified options from the base build in the **Modification and Customization Section**. Second, this build is highly modular; if a component fails or the user prefers a different alternative to a given module (*i.e.,* using an Arduino as opposed to a WAGO PLC), parts can easily be swapped or exchanged. This can be contrasted with commercial closed-source pumps and instruments where exchanges and repairs are often costly and time-consuming. Lastly, the control system described here is significantly cheaper per input than existing solutions, as discussed below.

### 2.4. Note on Associated Build Costs

This setup has a significantly reduced cost compared to commercial alternatives for the same number of independent input controls. **Table 1** shows the cost of hardware for **Build Option 1** (full build, 48 control lines), **Build Option 2** (modified programmable build, 8 control lines) and **Build Option 3** (fully manual, 12 control lines). The most expensive components are the solenoid valve arrays and WAGO fieldbus digital control modules, which scale directly with the number of independently-addressable control lines needed, as shown in **Table 1**. While the cost of these programmable setups can be significant ($4,455 to $10,380 dependent on number of inputs), commercial solutions cost between $1,500 (lower-end microfluidic syringe pumps) to $5,550 (high end pressure-driven pumps, *e.g.* Dolomite P-Pump^TM^) *per independent input.* Consequently, obtaining a similarly programmable pneumatic system for many inputs from commercial suppliers would be > 10X more expensive in the current market than the build described here.

**Table 1.** Cost of Hardware for the different Build Options. See Supplemental Bill of Material for itemized costs.

| Build Option | Cost of all associated hardware |
| --- | --- |
| #1: Full Programmable Build (48 control lines, 18 flow lines) | \$10,379.65 |
| #2: Abridged Programmable Build (8 control lines, 6 flow lines) | \$4,455.14 |
| #3: Manual Build (12 control lines, 18 flow lines) | \$2,110.50 |

### 2.5. Hardware Summary

- This build describes a pressure-driven pneumatic setup for single- and multi-layer microfluidic device operation. Devices can be operated manually or digitally, enabling remote control.
- The user can customize the number of flow inputs (up to 48) and the number of control inputs (up to 96) for their application.
- The associated software for digital valve control, Geppetto, includes a GUI for fast valve actuation and visualization of current valve states as well as an easy way to script experiments for automated control.
- The build description includes design files to be cut, printed, or machined, as well as overview instructions and step-by-step build guides to construct the setup in 5 different modules, a bill of materials and individual parts list by module, software operation instructions, and an example case for easy construction. These resources can also be found on our Github (https://github.com/FordyceLab/Open_Source_Pneumatics).
- The final setup is ∼48” × 24” including the control computer and can be mounted on an optical breadboard, lab bench, or any workspace with either screws or C-clamps.

## Design Files

### 3.3 Design Files Summary

The following design files are necessary to complete this build. Downloads of parts in editable file formats (*e.g.* AutoDesk 360 files) are available on the GitHub repository under the Designs folder.

**Table 2.** Design File Summary.

| Design file name | File type | Open source license | Location of the file |
| --- | --- | --- | --- |
| System Schematic | PDF | GPL | Available with the article |
| Bill of Materials, Detailed with Links | XLSX | GPL | Available with the article |
| Build Guides | PDF (multiple) | GPL | Available with the article |
| Base Board<br>(*backboard-full*) | PDF<br>DXF | GPL | Github: github.com/FordyceLab/Open_Source_Pneumatics |
| Flow Control Manifolds<br>(*stopcock-mount*) | STL | GPL | Github: github.com/FordyceLab/Open_Source_Pneumatics |
| Water Reservoir Stand<br>(*reservoir-stand*) | PDF<br>DXF<br>STL | GPL | Github: github.com/FordyceLab/Open_Source_Pneumatics |
| Control Water Reservoir<br>(*reservoir*) | PDF<br>DXF | GPL | Github: github.com/FordyceLab/Open_Source_Pneumatics |
| Geppetto<br>Geppetto CLI | Python code | MIT | Github: https://github.com/FordyceLab/geppetto<br>Github: https://github.com/FordyceLab/geppetto-cli |

### 3.4 Description of Design Files

#### Build Instructions

**System_schematic_pneumatics.pdf** – Schematic of all parts organized by build module

**Bill_of_Materials_pneumatics.xlxs** – Multi-sheet parts list, global and by-module (includes modification and custom build **Options 2** and **3**).

#### Build and Operation Step-by-Step Guides

**Module_1_Base_Build_Guide.pdf** – Guide for the Module 1 build

**Module_2_Filter_Assembly_Guide.pdf** – Guide for the Module 2 build

**Module_3_Valve_Manifolds_Guide.pdf** – Guide for the Module 3 build

**Module_4_Control_Loading_Guide.pdf** – Guide for the Module 4 build

**Module_5_WAGO_Guide.pdf** – Guide for the Module 5 build

**Operation_Start_Guide.pdf** – Instructions for setup initialization and first-time use.

**Device_Operation_Guide.pdf** – Instructions for device operation and experimental automation.

#### Laser Cut Parts

**Base Board: *backboard-full*** (PDF specifications; DXF for submission to 3^rd^ party) – 28” × 20” backboard to which most components are mounted. Fabrication: 1/4” aluminum or acrylic stock. Holes can be drilled or laser cut. Threaded holes hand-tapped as per specifications.

**Water Reservoir Stand: *reservoir-stand*** (PDF specifications; DXF for submission to 3^rd^ party; STL for 3D printing) – Mounting stand for control manifold water reservoirs. Fabrication: 1/4” aluminum or acrylic stock. Threaded holes hand-tapped as per specifications. Alternatively, can be ordered from local plastic suppliers/3^rd^ party, or 3D printed on printers with a sufficiently large workspace.

#### Optional Modified Laser Cut Parts

**Base Board Alternatives:**

> ***backboard-left*** (PDF specifications; DXF for submission to 3^rd^ party) – The left 10.5” of *backboard-full*. Its reduced size enables fabrication on laser cutters with smaller workspaces. It should be used with *backboard-right;* if this component is used, *backboard-full* is not needed.
>
> ***backboard-right*** (PDF specifications; DXF for submission to 3^rd^ party) – The right 17.5” of *backboard-full*. Same specifications as above.
>
> ***backboard-left-lc*** (PDF for laser cutter) – Variation of *backboard-left* specifically for direct conversion to a Universal Laser Systems file format (lines are 0.02 mm wide, [255,0,0] RGB). Please verify that the file format is compatible with your laser cutter.
>
> ***backboard-right-lc* (**PDF for laser cutter) – Variation of *backboard-right* specifically for direct conversion to a Universal Laser Systems file format (lines are 0.02 mm wide, [255,0,0] RGB).
>
> ***reservoir-stand-lc*** (PDF for laser cutter) – Variation of *reservoir-stand* specifically for direct conversion to a Universal Laser Systems file format (lines are 0.02 mm wide, [255,0,0] RGB).

#### Machined Parts

#### Control Water Reservoirs

***reservoir*** (PDF specifications; DXF for submission to 3^rd^ party) – Control manifold reservoirs used to hold water for control lines. Fabrication: 1” acrylic stock. Custom machined in-house or by 3^rd^ party.

#### Flow Control Manifold

***stopcock-mount*** (STL for printing) – Mount for 3 stopcocks that control flow line pressurization. Fabrication: ABS plastic or similar.

#### Open-Source Software

**Geppetto** (Github Repository) – Python pneumatic control software with Kivy GUI and optional Windows binary. A Read Me file and all code dependencies are included in the repository unless noted. Example files are included in the repository.

**Geppetto CLI** (Github Repository) – Python pneumatic control software for command line scripting (command line integration, CLI) to automate experiments. A Read Me file and all code dependencies are included in the repository. Example files are included in the repository.

## 4. Bill of Materials

### 4.3. Master Bill of Materials

The following is a global list of materials to be purchased before the build. Material quantities should be adjusted to the needs of your build. The main build (**Build Option 1**) used as an example in the following Build Guides is for 18 flow line inputs and 48 individually-addressable control line inputs.

Materials lists for modified **Build Options 2** and **3** are presented in the **Modification and Customization Section**.

**Table 3.**
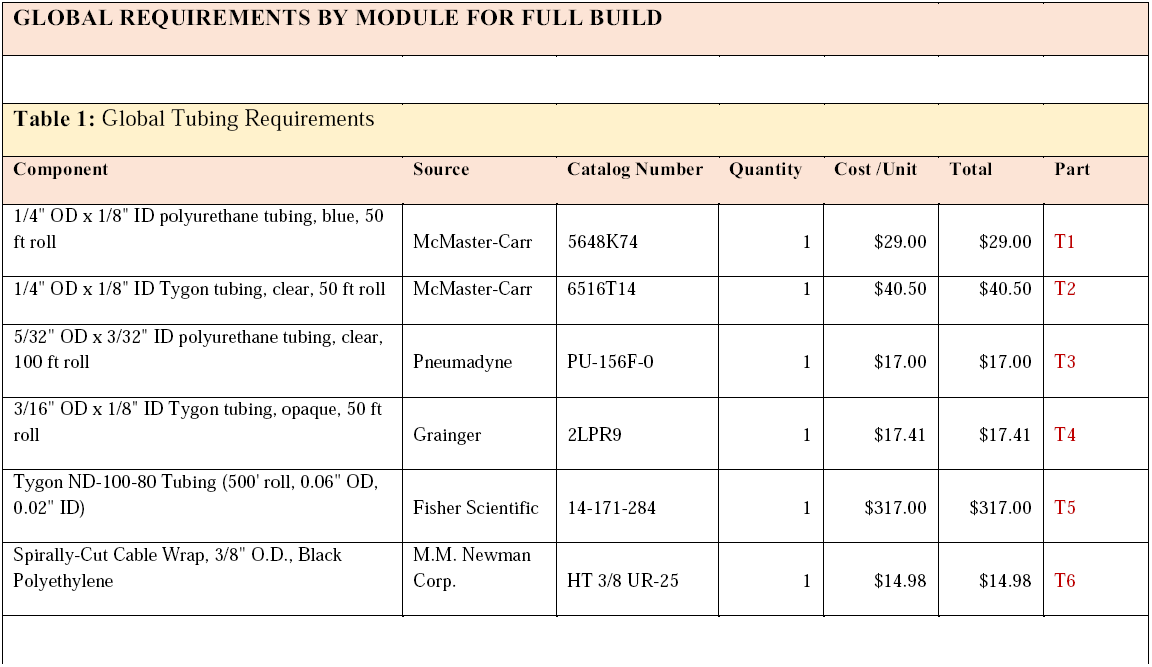
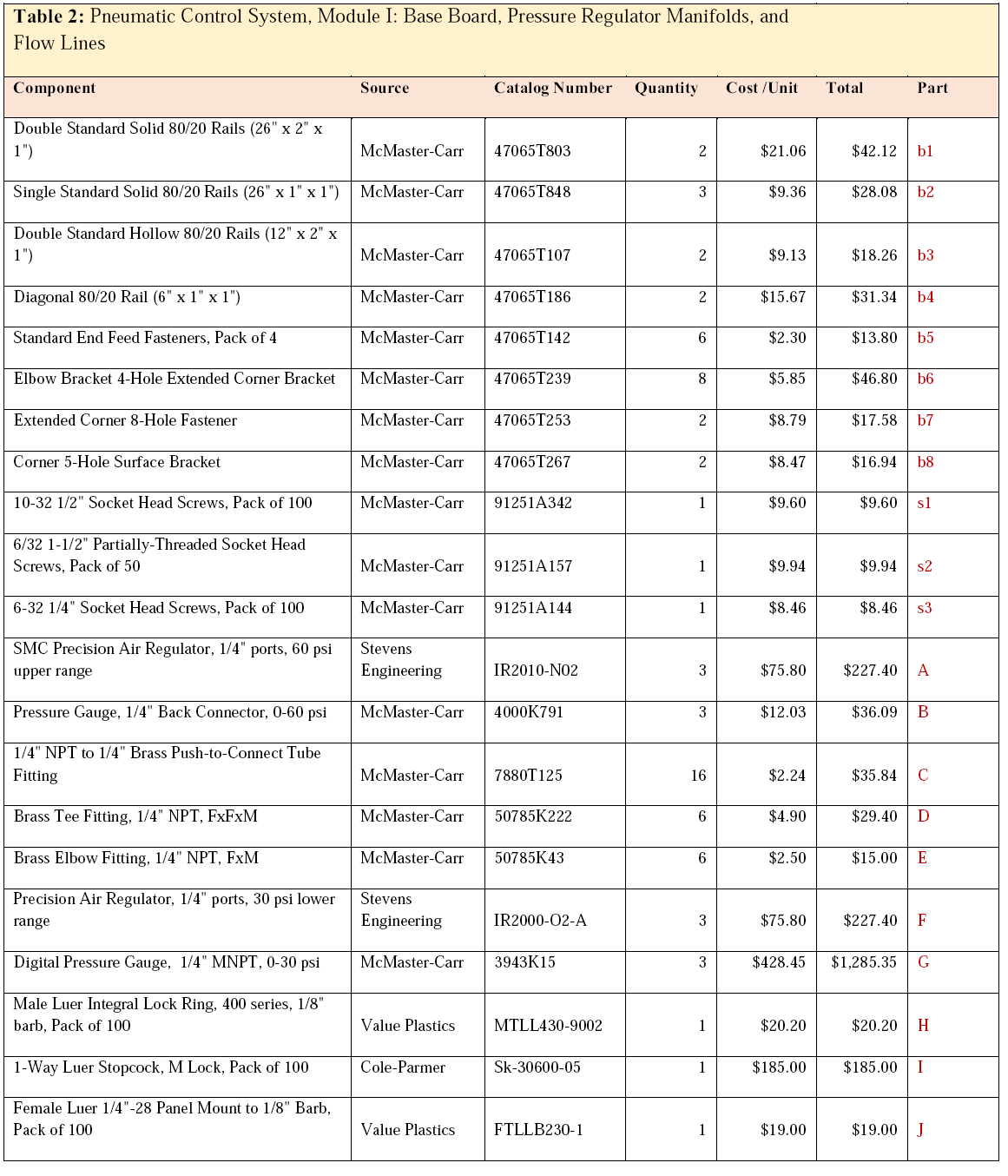
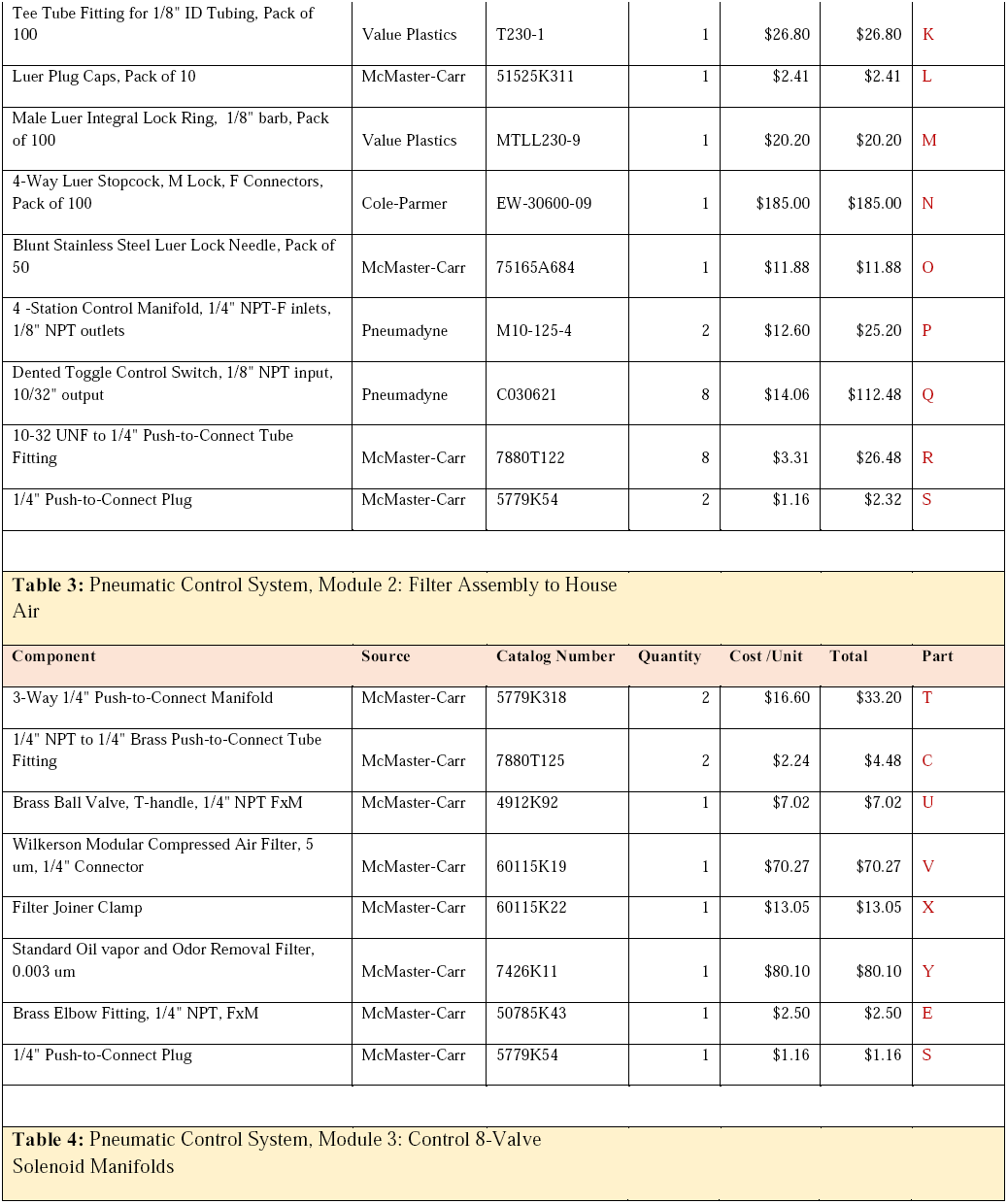
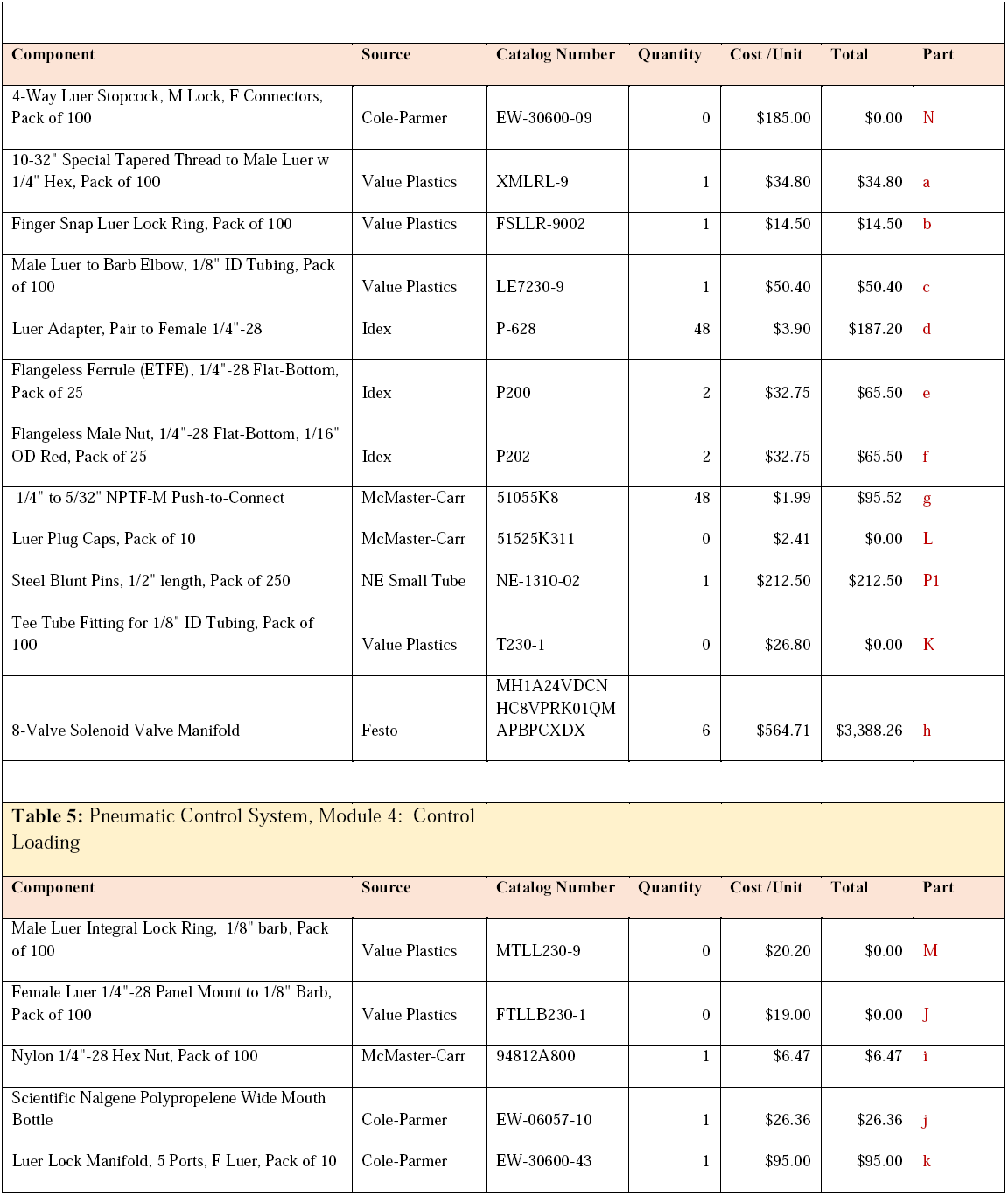
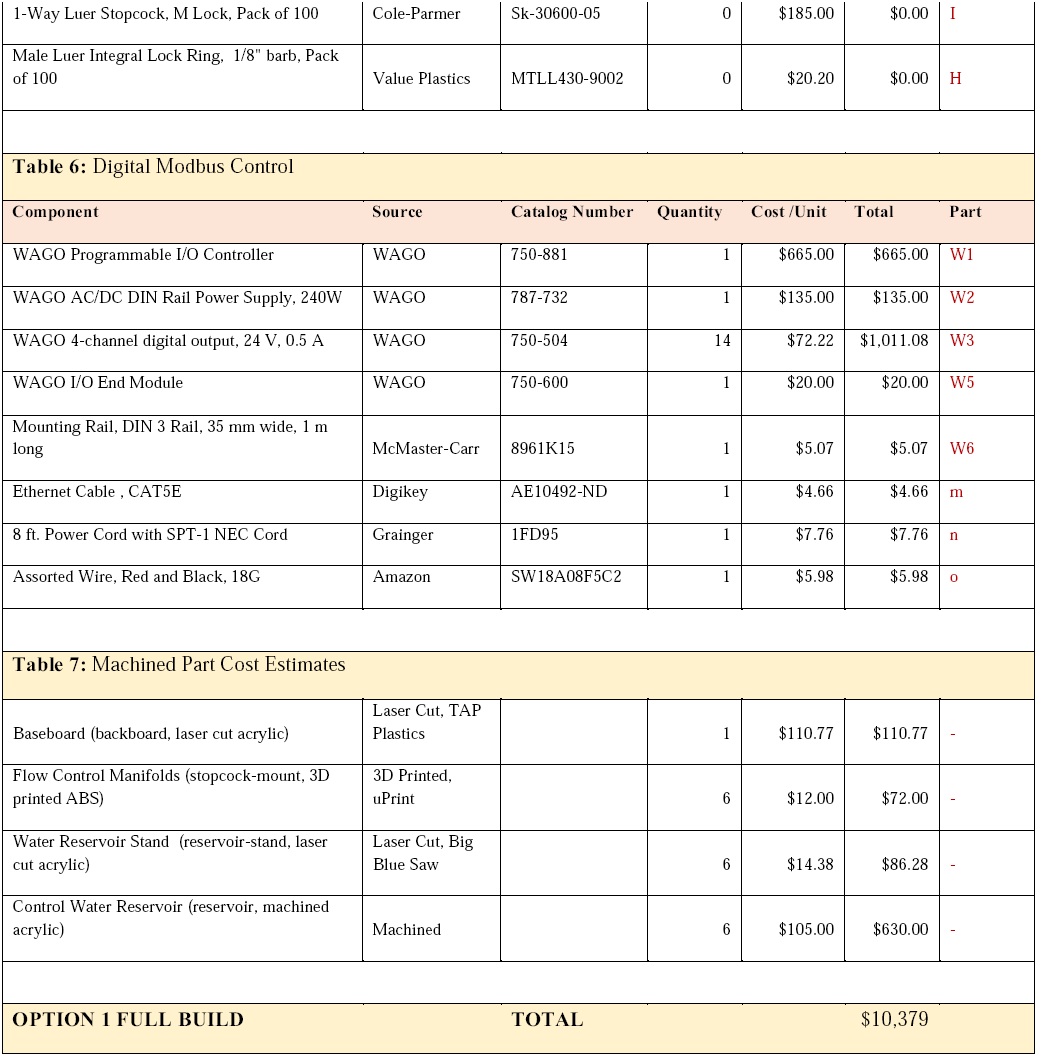
Full Build (Build Option 1) Bill of Materials by Module. Parts are purchased by Module. Any “0” quantity denotes that a multi-part pack has been previously purchased of sufficient size to provide all additional parts needed for the Module in question. Links to manufacturer sites are included in the **Bill_of_Materials_pneumatics.xlxs** supplemental file (**Supplement 2**).

A more detailed Bill of Materials with purchasing links and further information is available as **Supplement 2** and is updated often on our Github repository, available <u>here</u>.

## 5. Build Instructions

### 5.1. Overview of Multi-Component Build

This build takes place in 5 discrete modules that eventually interconnect. The following general build schematic describes all the parts and connections necessary to complete the build (**Figure 2**). For each module, we describe the module operation and provide a ‘close up’ version of the schematic compared to the final implementation of that assembly. Assembly details are presented step-by-step in detail in a series of 5 build guides in the **Supplemental Information** of this article. The end result will be a pressure-driven pneumatic setup for automated device control that is ready to use.

The following sections describe the details for the main **Build Option 1**. These sections can be used as examples for **Build Options 2** and **3** but will need to be adjusted accordingly to the schematics presented for those options, as shown at the end of this paper.

### 5.2. Module 1: Base Board, Pressure Regulator Manifolds, and Flow Lines

This module describes the main base build for the setup. Each build has air regulators to control the pressures used to drive manual flow and valve lines. As different on-chip flow inputs may require different pressures, the board contains several different ‘flow’ regulators to control pressures to the flow line manifolds mounted on the base board (right side regulators). In the main build, 3 regulators control 18 flow lines as shown in **Figure 2**. Digital gauges provide highly precise low-range (0-20 psi) flow control and are powered by the WAGO modus controller described in **Module 5**. The manual build, **Build Option 3**, replaces these digital gauges with low-range (0-10 psi) analog gauges for a cost-effective alternative, with some sacrifice to measurement precision.

For automated builds (**Builds 1** and **2**), regulators also supply pressure for the banks of control solenoid valve manifolds that ultimately pressurize the control lines and actuate on-chip microfluidic valves (left side upper regulators). In the main build, 2 regulators control pressurization of the control solenoid manifolds and modulate their pressurization via mounted switchboxes (left side). These switchboxes connect directly to the 8-valve manifolds presented in **Module 3**. The regulators used here can be relatively imprecise as long as they provide pressure regulation in the range needed to close microfluidic devices (0-60 psi). Manual versions are sufficient. Automated builds further have an integrated option to load and refresh the control manifold valve banks with water easily. In the main build, a regulator (left side bottom regulator) is dedicated to this task and connected to the control loading assembly in **Module 4**. These components are mounted to a single base plate for convenient adjustment and connection between modules. The build guide provides instructions on how to mechanically connect all components to the base plate and pneumatically connect components to one another.

**Figure 3**. (a) Module 1 schematic. (b) Finalized implementation of Module 1.

**Figure 3.**
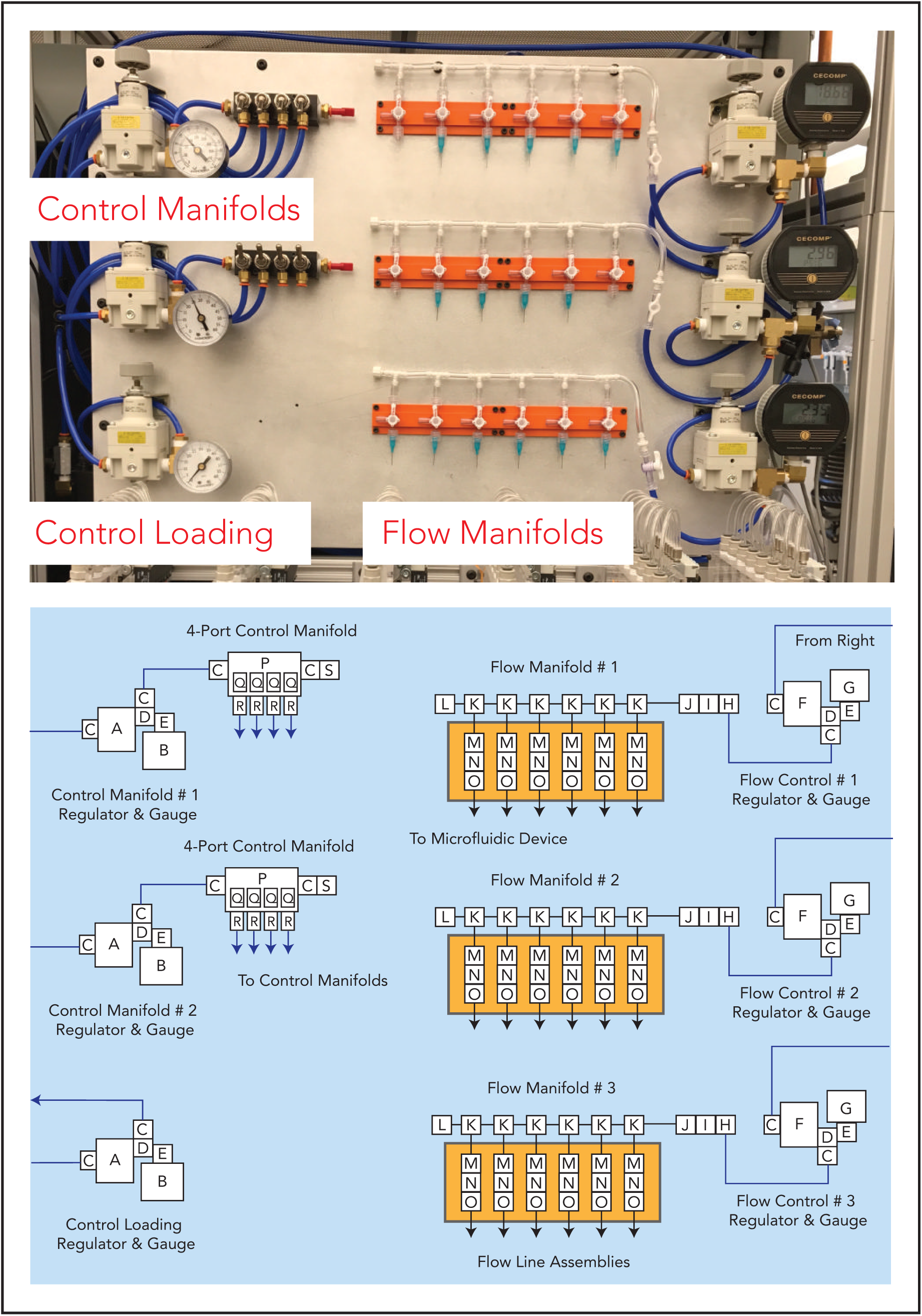
Module 1. **(a)** Module 1 schematic. **(b)** Finalized implementation of Module 1.

To build this module, follow the **Module_1_Base_Build_Guide.pdf** (Supplement 3).

### 5.3. Module 2: Filter Assembly

This module inserts a filter assembly between the compressed air source and the regulators on the Base Board (**Module 1**) to keep oil or small particulates in the pressurized air from contaminating the regulators and on-chip valves in critical experiments (*e.g.* long-term cell culture or on-chip enzymatic assays). This assembly can be replaced by cheaper disposable filters if desired.

**Figure 4**. (a) Module 2 schematic. (b) Finalized implementation of Module 2.

**Figure 4.**
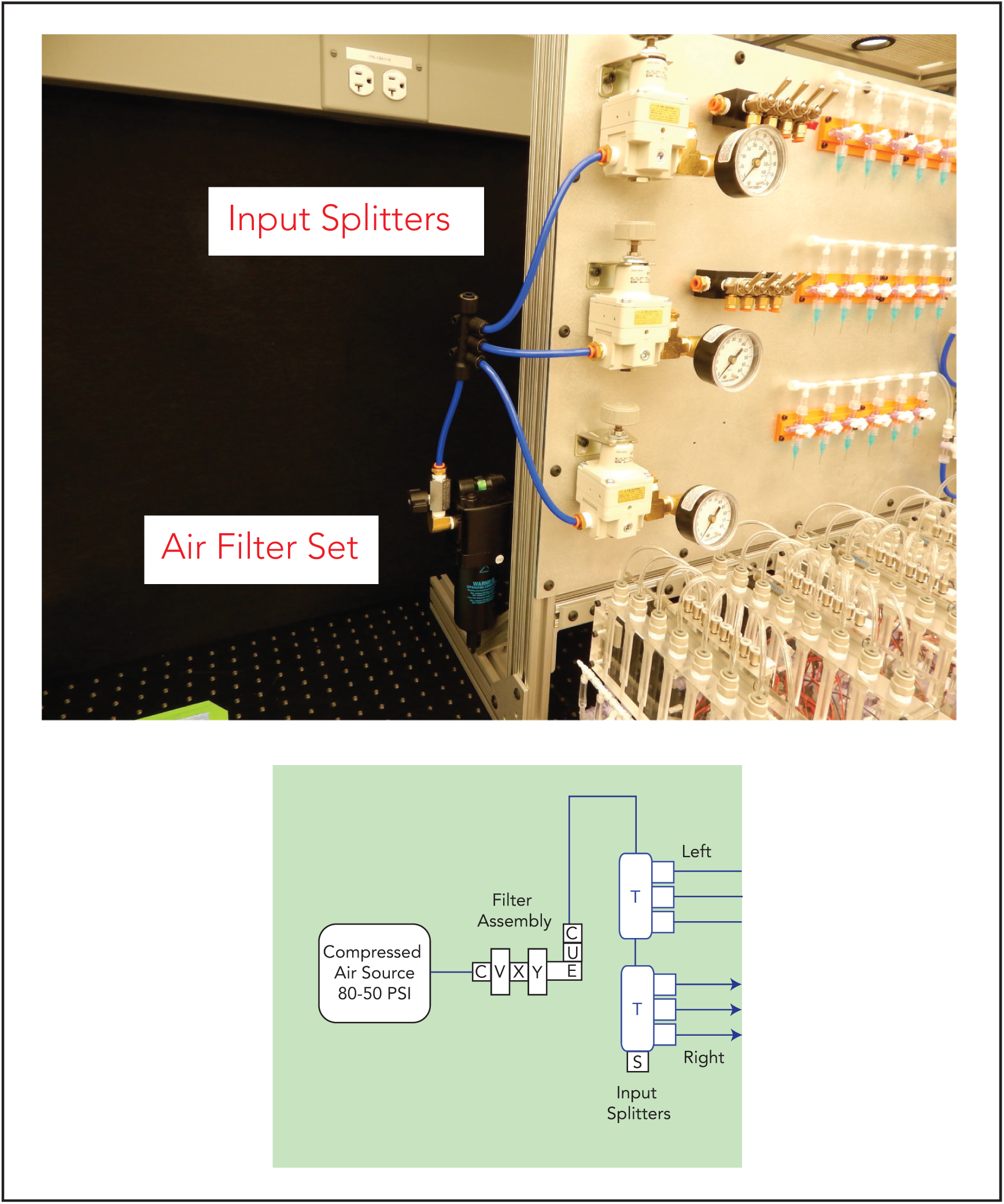
Module 2. **(a)** Module 2 schematic. **(b)** Finalized implementation of Module 2.

To build this module, follow the **Module_2_Filter_Assembly_Guide.pdf** (Supplement 4).

### 5.4. Module 3: Control 8-Valve Solenoid Manifolds

This module creates the hardware necessary for automated valve actuation. The main build has 6 sets of Control Valve Manifolds that actuate on-chip valves by pressurizing water-filled Tygon control lines connected to on-chip control channels. Each Control Valve Manifold has 8 water reservoirs connected to 8 solenoid valves with independent actuation. Each solenoid valve is individually driven by a 24V signal supplied by the fieldbus controller of **Module 5**. When actuated, each solenoid pressurizes a water reservoir (pressure sent from the Base Board – **Module 1**) and its connected control line input to the microfluidic device to close dead-end valves. The water reservoirs also have a loading function (via a separate fluidic circuit activated by toggling the manifold stopcocks) so that water can be refreshed via **Module 4**.

Fast valve pressurization is often critical for many microfluidic assays including on-chip biochemical assays, droplet generation, microfluidic sorting and precise fluidic dosing using on-chip pumps [20,22-24,35]. The Festo 24V solenoid valves used here, while expensive, offer superior response times essential for fast valve actuation as well as reliable state switching with long lifetime (>5 years operation). The response times of these solenoid valves is ∼ 4 ms response from pulse. Full fluidic valve actuation on-chip from the digital pulse is on the order of 7-20 ms, dependent on valve and device geometry. The three-way, normally-open solenoid valves in this setup toggle between atmospheric pressure and compressed air as set by the regulators of **Module 1,** allowing lines to be vented to atmosphere when not in use (two-way valves lack the ability to vent lines or depressurize lines to open valves). The use of normally-open solenoid valves allows on-chip valves to return to a safe ‘closed’ position in the event of an unexpected software crash or power interruption. In addition, these valves are additional very low profile, easily allowing parallelization as demonstrated in this build.

To customize the build, control lines can be filled with oil or buffer instead of water to reduce osmotic water transfer between control and flow lines. Three-way solenoid valve arrays can also be connected to other air sources to toggle between different states (*e.g.* vacuum and compressed air, vacuum and atmospheric pressure – instead of atmospheric pressure and compressed air), if needed.

**Figure 5**. (a) Module 3 schematic. (b) Finalized implementation of Module 3.

**Figure 5.**
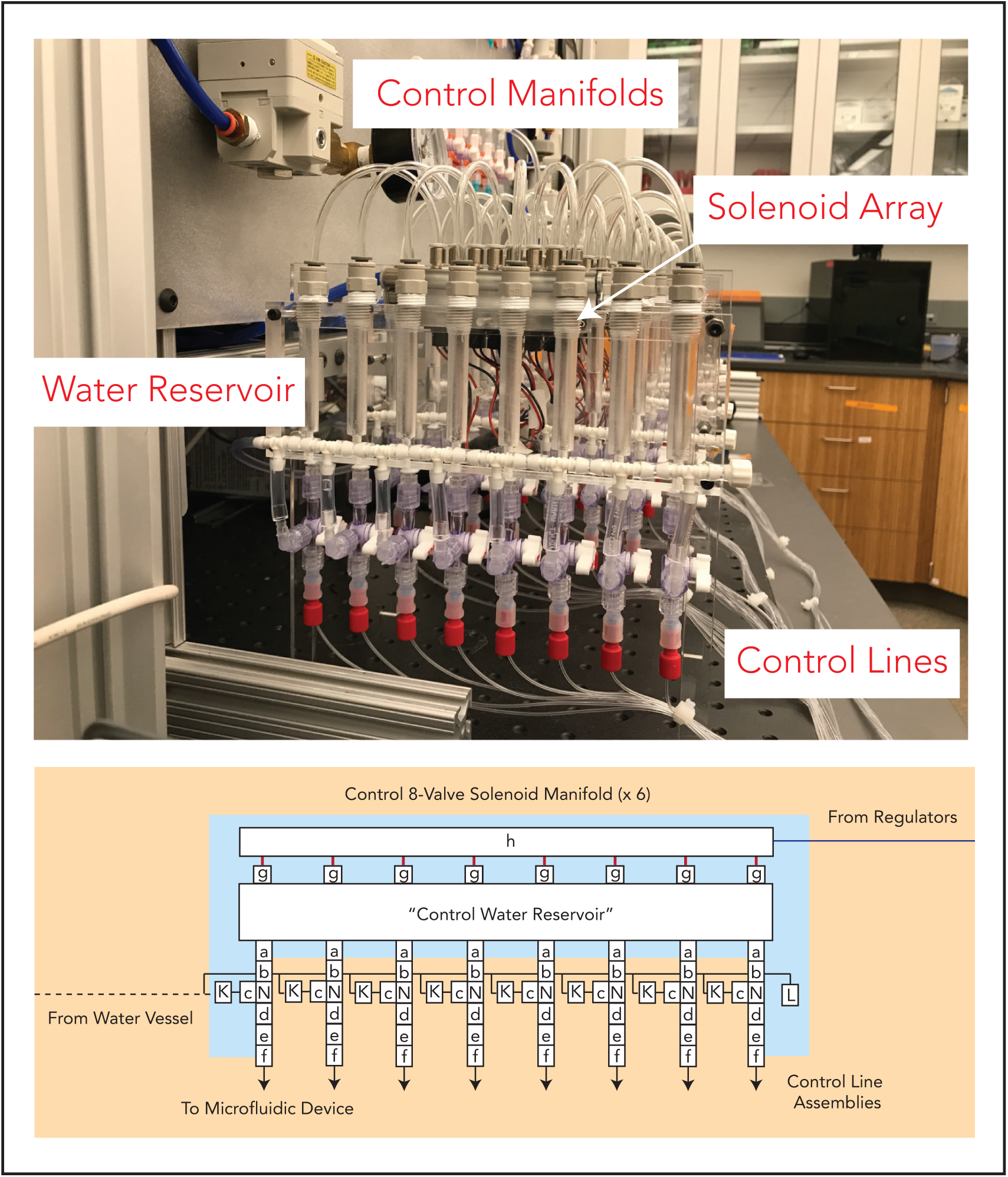
Module 3. **(a)** Module 3 schematic. **(b)** Finalized implementation of Module 3.

To build this module, follow the **Module_3_Valve_Manifolds_Guide.pdf** (Supplement 5).

### 5.5. Module 4: Control Reservoir Loading Assembly

Maintaining water in the water reservoirs is essential for properly pressurizing control lines as air within the lines can create uneven closing times across the devices and nucleate bubbles in the flow channel in long-term experiments. This module builds a pressure vessel and associated tubing for loading water into the water reservoirs of the Control Valve Manifolds from **Module 3** to prevent control line drying. While loading the water reservoirs can be completed without this pressure vessel assembly (*e.g.* using tubing), this assembly simplifies the water loading process. Water is loaded by pressurizing the water vessel (driven from a regulator in **Module 1**) and switching the relevant manifold stopcock so that water flows into the manifold. Further details can be found in the **Device Operation Guide**.

**Figure 6**. (a) Module 4 schematic. (b) Finalized implementation of Module 4.

**Figure 6.**
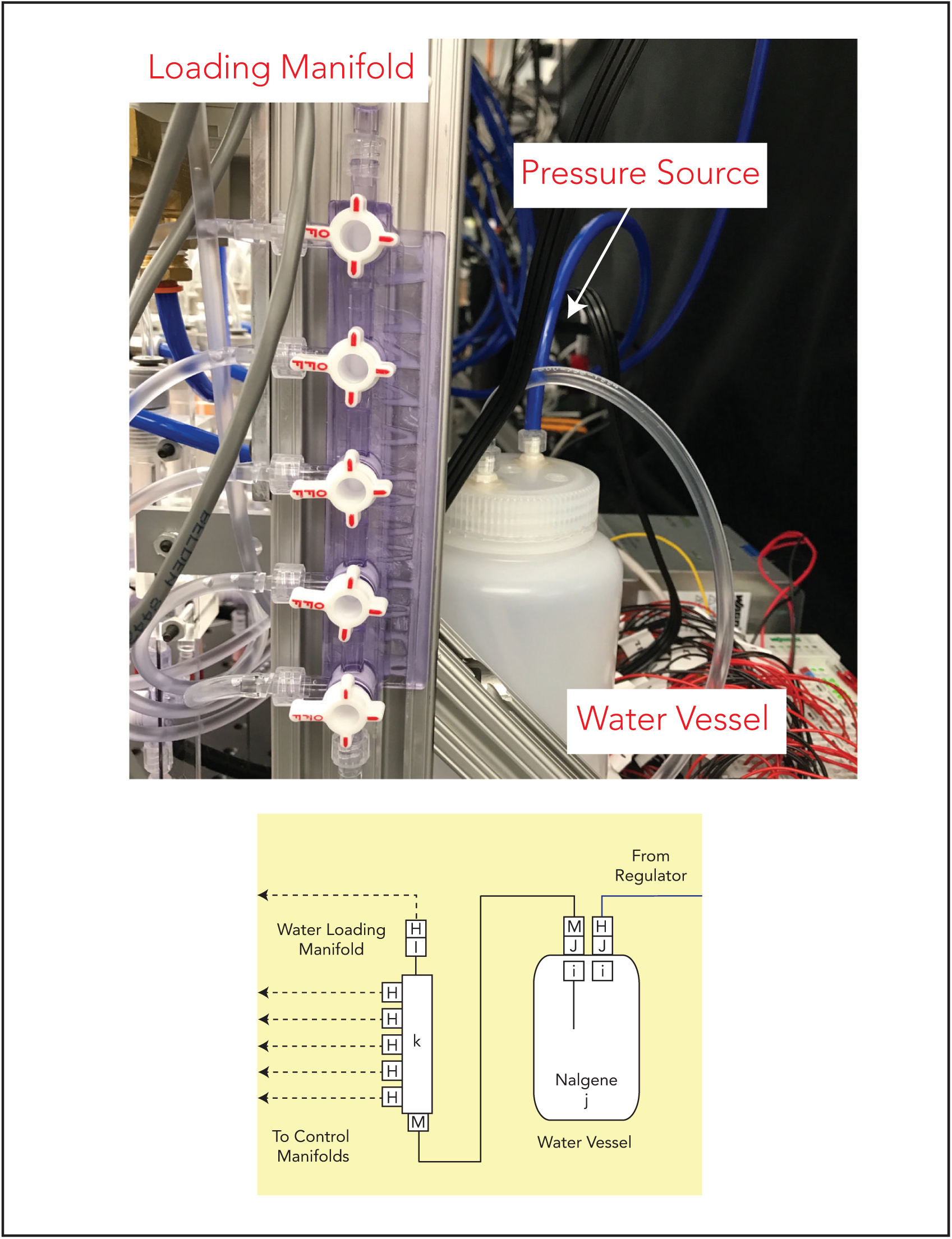
Module 4. **(a)** Module 4 schematic. **(b)** Finalized implementation of Module 4.

To build this module, follow the **Module_4_Control_Loading_Guide.pdf** (Supplement 6).

### 5.6. Module 5: Digital Modbus Control

This module adds computer-based control of valve actuation. A WAGO programmable logic controller (“Fieldbus controller”) is used to send 24V pulses to actuate the solenoid valves of **Module 3** individually, as well as power the digital gauges of **Module 1** (*note:* these gauges can be powered without the WAGO, but the WAGO provides a convenient integration of all electronic components in this context). The WAGO controller accepts a wide variety of input/output modules (“brick modules”) to create flexible and scalable control systems. The build presented here includes a power supply module for global power and ground, the WAGO controller CPU module with Ethernet connectivity, a series of 24V digital output (DO) modules for controlling valve states, and a 5V output module for powering auxiliary devices (*e.g.* digital gauges or hardware requiring TTL inputs). A computer can communicate with the WAGO fieldbus controller over Ethernet using a standard protocol called MODBUS. These communications are abstracted from the user using our Geppetto GUI or CLI software. Full wiring and computer setup details are reviewed in the Build Guide for this module.

For project-specific customization, it should be noted that the 4-way DO modules we have chosen send out digital signals that switch between 0V (float) and 24V (high-side control). Modules for more complex tasks such as digital in (DI) or mixed DO/DI modules can be swapped in or out depending on your needs (*e.g.,* for additional equipment control). Alternatives to the WAGO controller (such as Arduino PLCs, presented by White and colleagues in this issue [ref]) can be used but current and voltage compatibilities with the Festo solenoids should be checked before wiring.

**Table 4.** WAGO global settings for this implementation.

| WAGO Global Setting | File type |
| --- | --- |
| WAGO Ethernet Dip Switch | 198.168.1.3 |
| Connection Watchdog Timer (Ethernet) | 0 (timer off) |

**Figure 7**. (a) Module 5 schematic. (b) Finalized implementation of Module 7.

**Figure 7.**
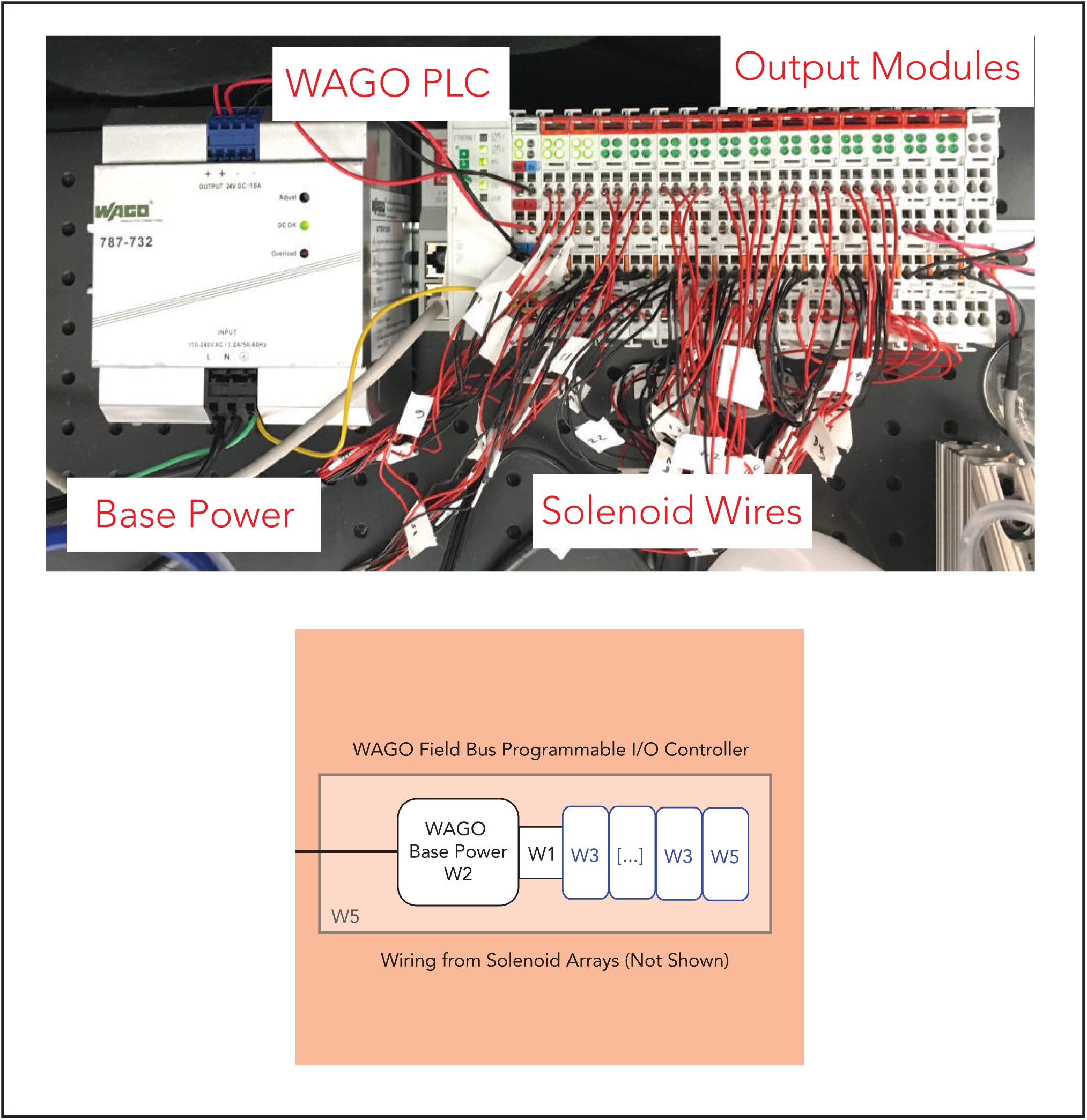
Module 5. **(a)** Module 5 schematic. **(b)** Finalized implementation of Module 5.

To build this module, follow the **Module_5_WAGO_Guide.pdf** (Supplement 7).

## 6. Modification and Customization

The pneumatic setup outlined here can easily be customized for your laboratory’s needs. Each component is highly modular, enabling easy part exchanges. Our Build Guides contain useful advice on our own trials using different part alternatives, when applicable. To demonstrate the ability to modify the base build, we present two additional build options with accompanying schematics. To build these options, simply use the previous Build Guides as needed for each part; component installation is exactly the same. The fully manual build, **Build Option 3**, does not use **Modules 3, 4,** or **5.**

> **Option 2:** An abridged build for 6 flow lines and 8 individually addressable solenoid valves to control 8 control lines that fully automated, programmed and remotely operated for simpler devices

**Figure 8**. Build Option 2 Schematic.

**Figure 8.**
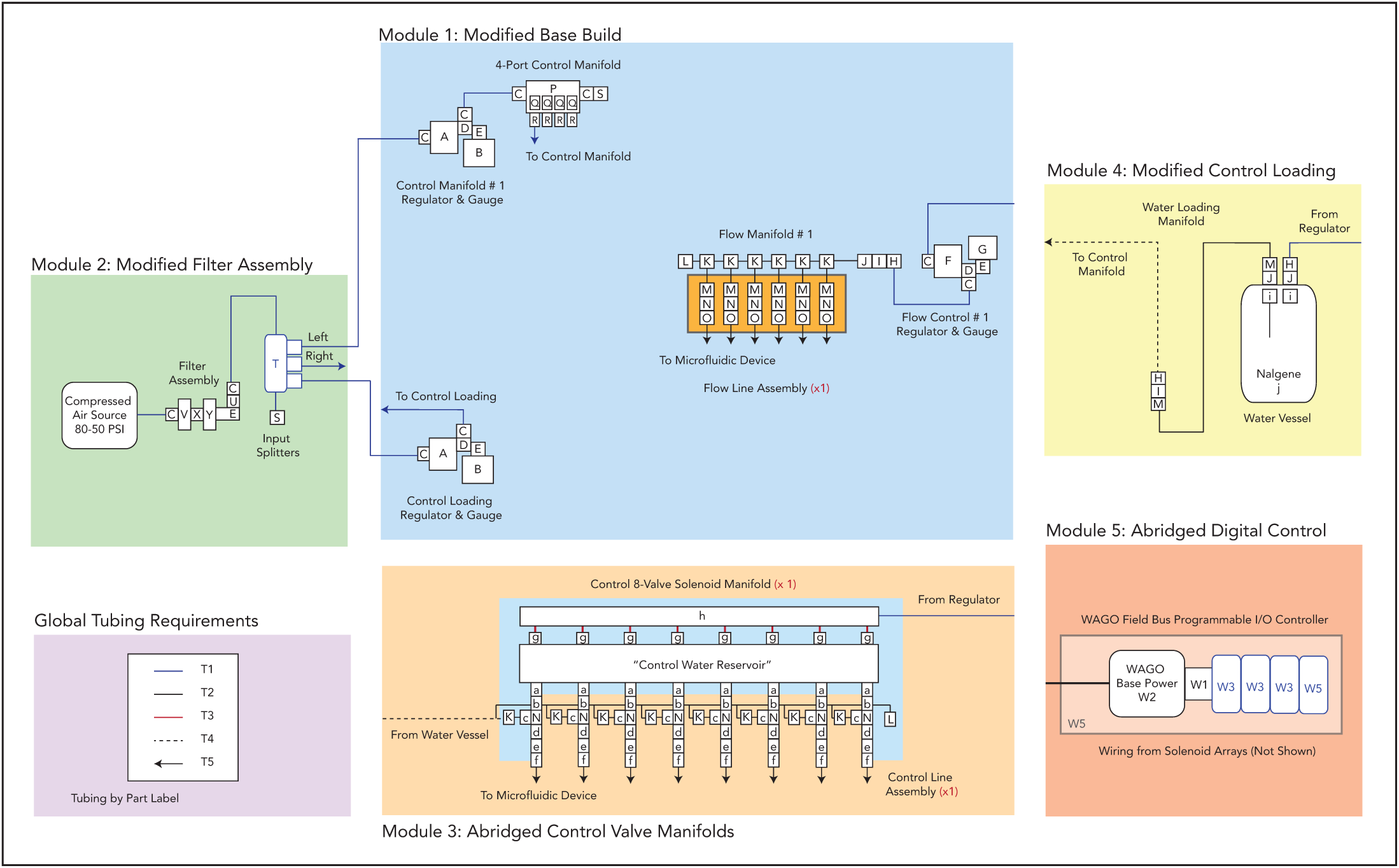
Build Option 2 schematic.

To build this option, follow the **Option 2 Build Materials List** (Supplement 2, Bill of Materials Sheet 2).

> **Option 3:** A fully manual build for 18 flow lines and 12 manually addressable control lines without automation capabilities which can be used to save costs

**Figure 9**. Build Option 3 Schematic.

**Figure 9.**
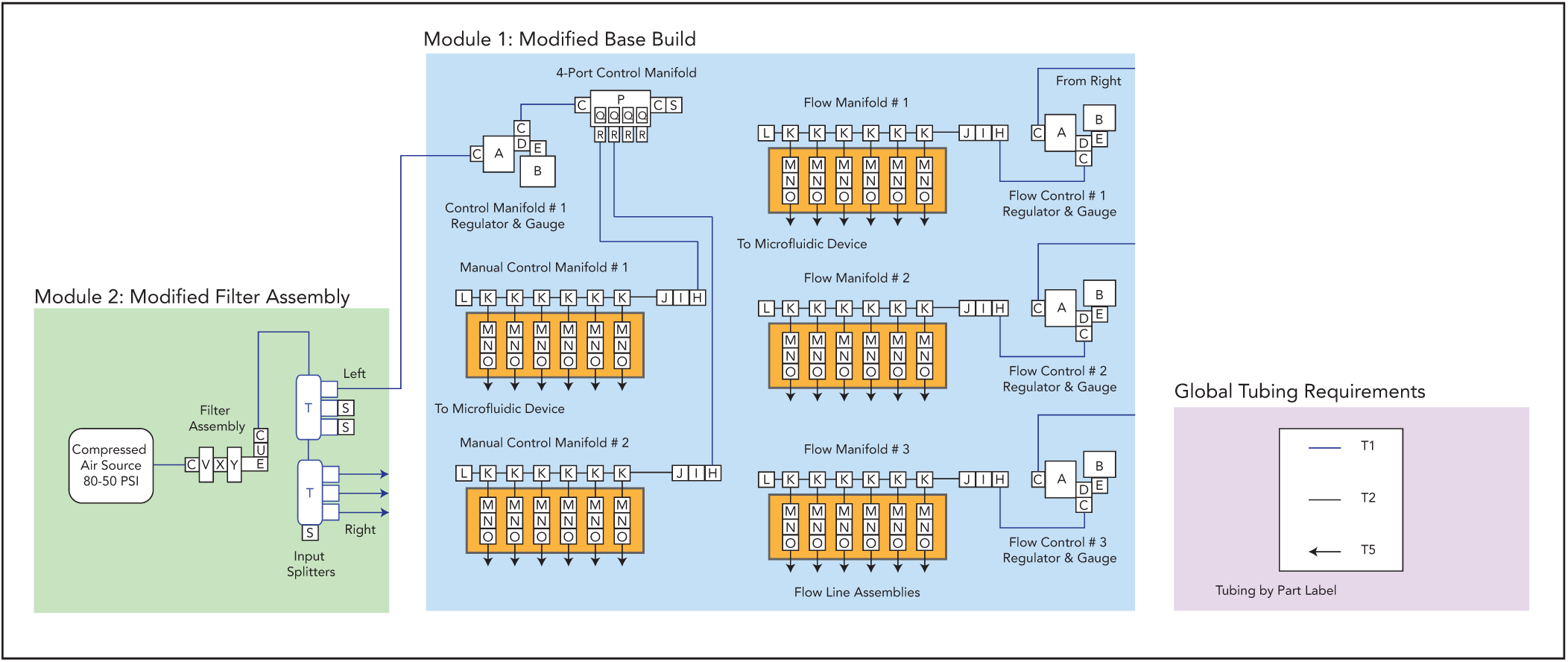
Build Option 3 schematic.

To build this option, follow the **Option 3 Build Materials List** (Supplement 2, Bill of Materials Sheet 3).

## 7. Operation Instructions

After completing the build, it is now time to begin the exciting process of testing and operating the first microfluidic device with your new setup! Before beginning, follow the initialization guide **Operation_Start_Guide.pdf** (Supplement 8) to pressurize and check for leaks in the system.

### 7.1. Geppetto: Python Operation and Scripting

We recommend operating your microfluidic device using our pneumatic control software, which enables valve state visualization and control as well as scripting. Alternatively, users can write their own MODBUS implementation for talking to the WAGO, if desired. Geppetto is a Python-based pneumatic control software built using the <u>Kivy graphical user interface framework</u> for GUI development.

Geppetto’s GUI supports nearly any user-defined control layout, with the ability to change the background image and the placement of the control buttons to match any user-designed device using a simple <u>YAML</u> config file.

To customize Gepetto for your own microfluidic device, use the Read Me under our Github release available at https://github.com/FordyceLab/geppetto. If you are unfamiliar with Python, an excellent tutorial can be found <u>here</u> on how to install and operate Python on your machine.

**Figure 10**. Example Geppetto GUI for valve state visualization of a microfluidic device. Shown here is the custom GUI configured for the MITOMI demo device (see Section 9).

**Figure 10.**
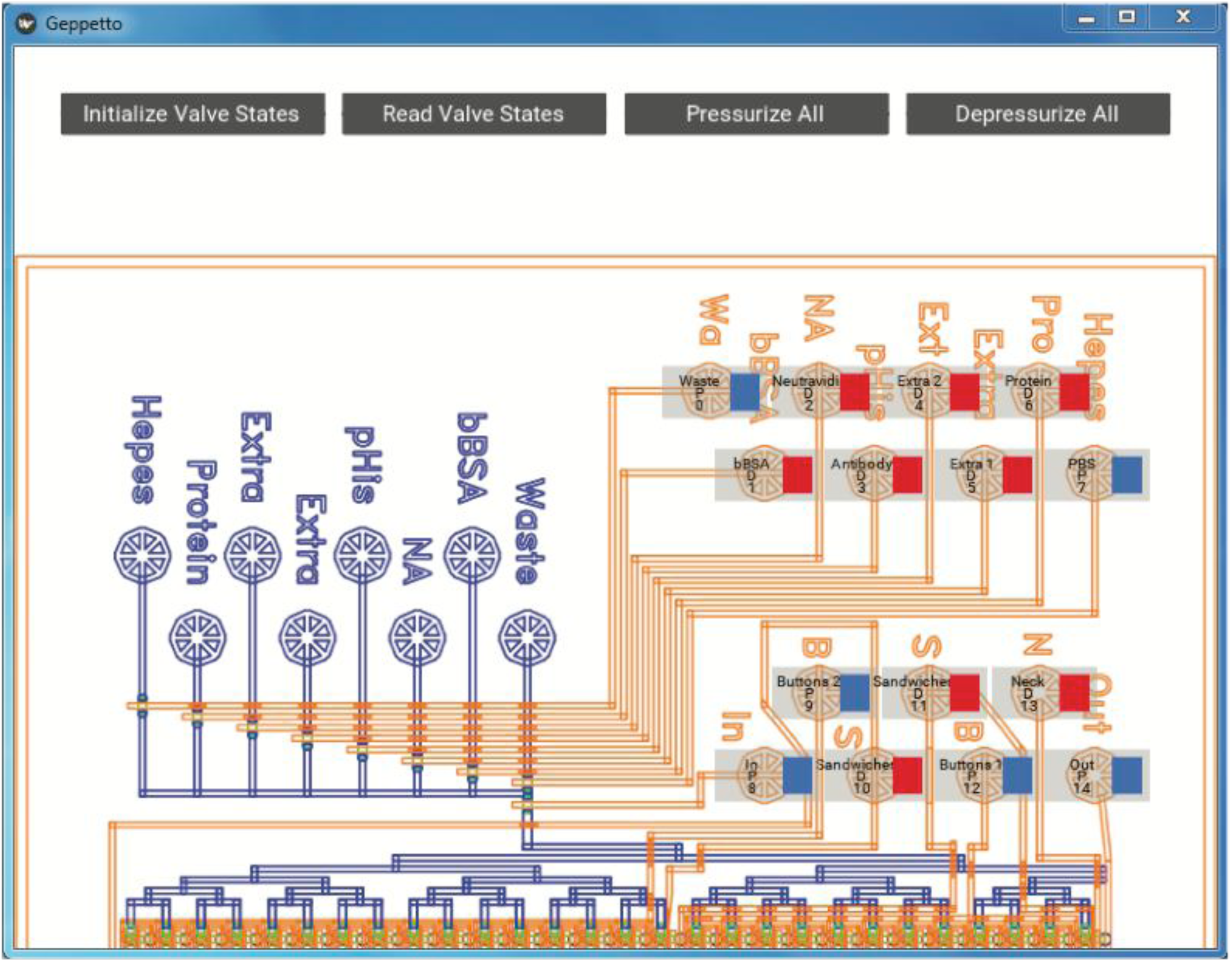
Example GUI for valve state visualization of a microfluidic device. Shown here is the custom GUI configured for the MITOMI demo device (see Section 9).

After installing Geppetto, load an image of the device design according to instructions in the Read Me and adjust valve locations to the correct positions in the image. The final image should look similar to the following (**Figure 10**).

### 7.2. Operating a Chip

To operate a microfluidic device, follow the steps **Device_Operation_Guide.pdf** (Supplement 9).

To summarize:

1. Connect the control lines and flow lines to the device.
2. Pressurize the control and flow regulators to the desired pressures and toggle the control manifold switches to send pressure into the control manifolds.
3. Use Geppetto to actuate the valve states.

We previously published a Jove video protocol of a device run that may be useful for visualization of this process [32].

### 7.3. Automating Microfluidic Experiments

Geppetto also allows scripting of valve actuation events and wait times to automate microfluidic experiments. Scripting occurs in the command line using the Geppetto CLI program (https://github.com/FordyceLab/geppetto-cli). Use the Read Me to set up your first scripting experiment. Example files (*e.g.* the script for our example experiment below) can be found under the Scripts folder.

## 8. Validation and Characterization

The pneumatics control setup described here has been used in our laboratory for multiple studies, including the microfluidic generation of hydrogel microparticles [32], microfluidic synthesis of lanthanide-encoded beads [24], and high-throughput biochemical affinity measurements on-chip [35].

Here, we describe an example microfluidic experiment using the full pneumatic build, Geppetto CLI, and a microfluidic device, MITOMI (as described previously [19,20,35]) for high-throughput affinity measurements.

### 8.1. Confirmation of Multi-valve Pressurization and State Change

**Figure 11** shows a brightfield image of multiple valve opening and closing events on chip, with valve state controlled via the Geppetto GUI. **Supplement 10** is a video of valve toggling events, demonstrating individually-addressable valve control.

**Figure 11.**
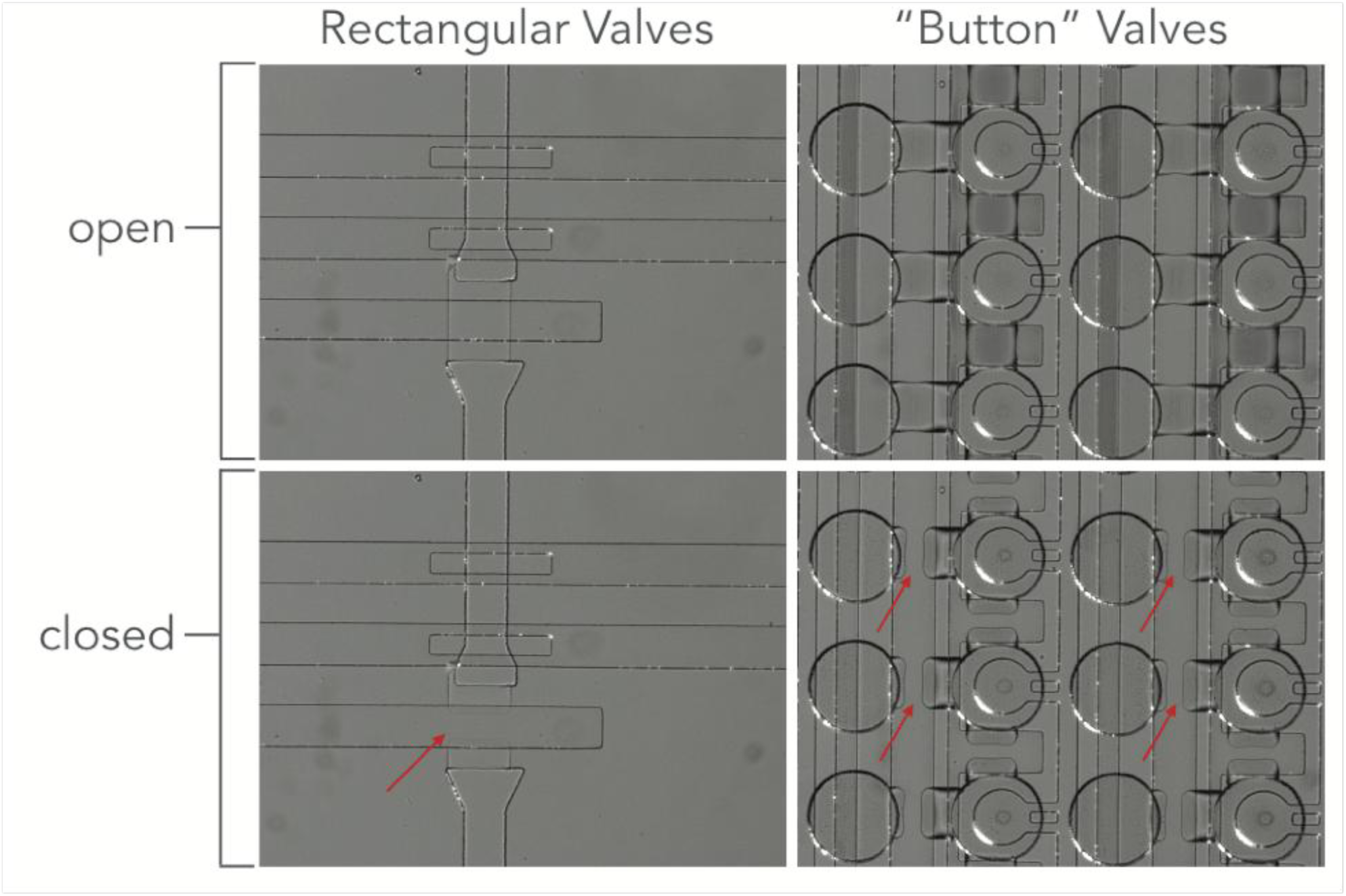
Example valve actuation. Valve open and closed (red arrows) states for 2 types of valve geometries under 25 psi control on a MITOMI device.

**Figure 11**. (a) Valve closed state under 25 psi control on a MITOMI device. (b) Valve open state.

### 8.2. Characterization of Valve Closing Pressures and Response Times for Multiple Device Geometries

We have characterized required microfluidic valve control pressures across a wide range of device geometries. “Push-down” devices (control layer on top) fabricated via standard soft lithography techniques tend to operate at 15-20 psi for complete valve closure. “Push-up” devices (control layer on bottom) can tolerate higher pressures (*e.g.* these devices are more resistant to delamination) and operate from 10-40 psi (or higher). For a given channel geometry, push-down valves require higher pressures to close than push-up valves [2,10,11]. The appropriate valve closing pressures can be determined by the user for new device geometries by introducing using beads or food dye into the flow channel, varying input pressures, and visualizing whether all the fluid is completely excluded beneath the valve.

### 8.3. Automating a Microfluidic Experiment

To demonstrate the automation potential of the setup, we scripted an experiment in Geppetto to carry out successful surface patterning of a MITOMI device. MITOMI is a microfluidic device that has been employed for many high-throughput biochemical measurements, most notably for assessing thousands of transcription factor (TF) binding affinities to DNA sequences printed on a glass slide [19,20,35]. MITOMI devices contain hundreds to thousands of individual pairs of chambers with multiple types of valves (*e.g.* a neck valve for connecting chamber pairs, button valves for patterning specific areas in a larger chamber, and sandwich valves for isolating chambers from one another) for performing biochemical affinity measurements.

In our experiment, we performed a simple multi-step surface functionalization to pattern anti-eGFP in a small circular area in each chamber pair using the ‘button’ valves of the device, which protect the glass surface from interacting with introduced solutions when activated. We automated the experiment using scripting in Geppetto CLI. Our script (**Supplement 11**) outlines the valve actuation events and flow times. Further details on this experiment program can be found <u>here</u>.

In particular, we created a patterned circle of anti-eGFP antibodies on the glass slide surface in the “push-down” device via the following steps: (1) open button valve, (2) introduce biotinylated BSA, (3) wash, (4) introduce neutravidin, (5) wash, (6) close button, (7) passivate all surfaces *except those beneath the button* by introducing more biotinylated BSA, (8) wash, (9) open button, (10) introduce biotinylated anti-eGFP (which is recruited to neutravidin-coated surfaces beneath the button), and (11) wash. Device preparation and setup for this experiment is described in detail as an example in the **Device_Operation_Guide.pdf** (**Supplement 9**).

We then tested if the patterning had been successful by flowing eGFP protein over the device (button valves open) followed by washing and then visualizing the resultant fluorescence in the eGFP channel. Surface patterning and immobilization is shown in **Figure 13**, demonstrating that valve actuation and automated control of sample introduction during the experiment were successfully implemented.

**Figure 13.**
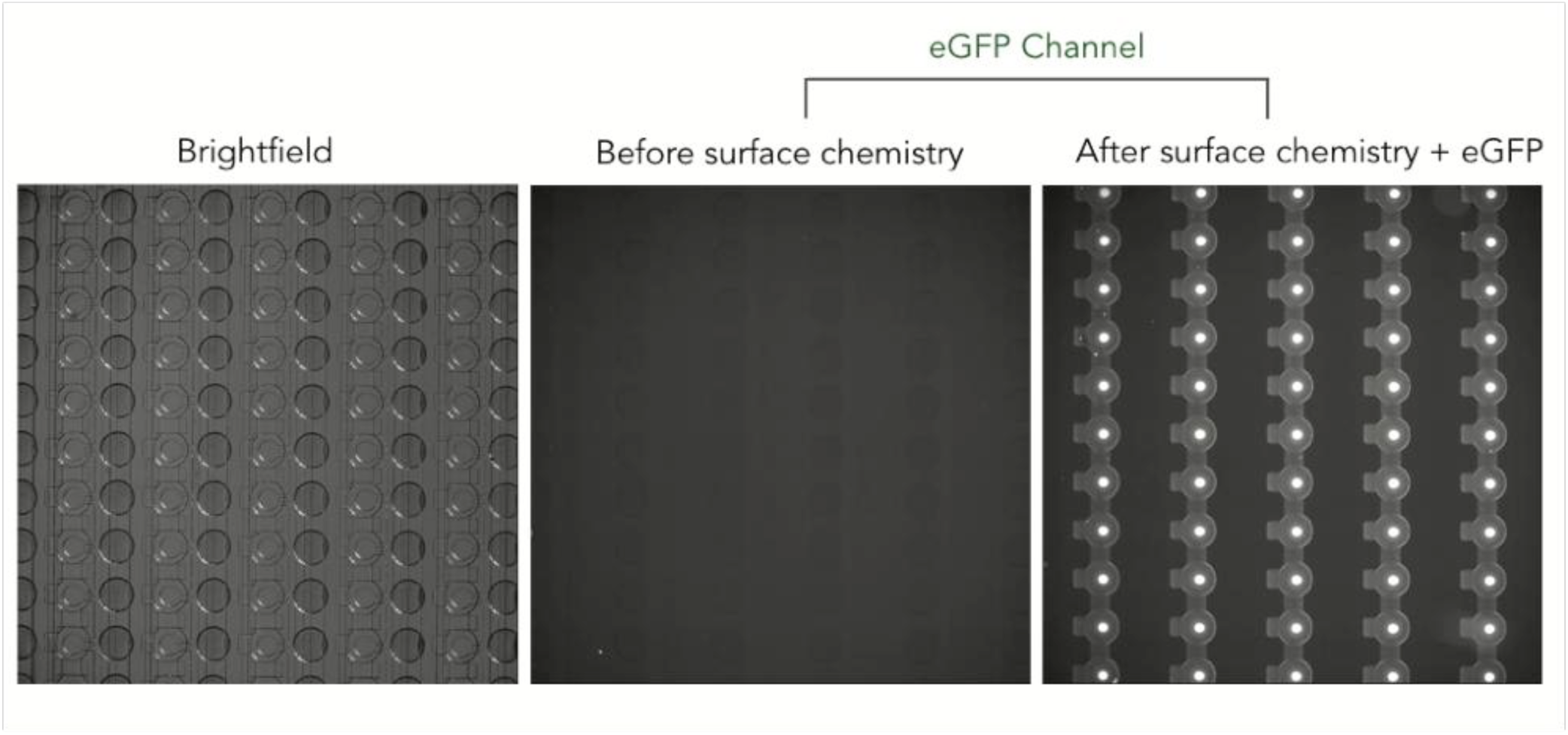
Experimental results showing an increase in eGFP only under the button area resulting from precise valve actuation events for successful patterning.

**Figure 12**. Image of a fully prepped MITOMI device with control lines and flow lines in operation.

**Figure 12.**
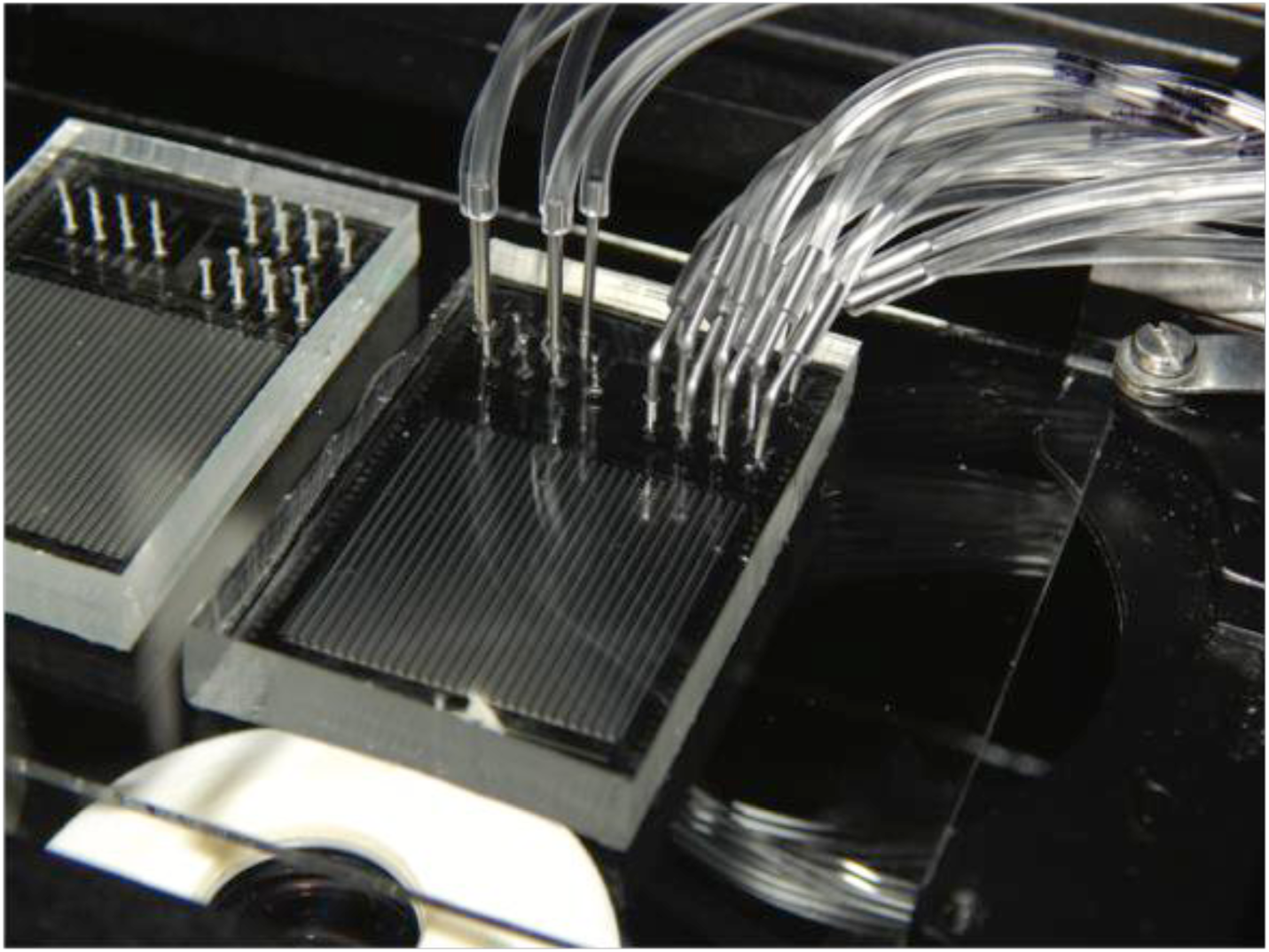
Image of a fully prepped MITOMI device with control lines and flow lines in operation.

**Figure 13**. Experimental results showing an increase in eGFP under only the button area resulting from precise valve-actuation events for successful patterning.

## 9. Discussion

In this report, we provide instructions on how to build a pressure-driven pneumatic setup for universal microfluidic device operation and automated control. This system is capable of driving 18 flow inputs and 48 individually-addressable control line inputs on a microfluidic chip. The setup build is easily adjustable, scalable, and customizable to a user’s needs while being cost-effective compared to similar commercial solutions. To demonstrate this modularity, we have presented two additional build options, (1) an abridged programmable setup with fewer control lines and (2) a fully manual setup. With the completion of the build, any user should be able to operate and automate a wide range of microfluidic devices and experiments. This open-source hardware and software build should enable researchers from all fields to take advantage of microfluidic technologies and easily operate their own devices.

## 10. Acknowledgements

The authors acknowledge the Fordyce Lab members, in particular Bianca Cruz, for their helpful comments and feedback on the build revisions and manuscript. This work was supported by NIH/NIGMS grants R00GM09984804, DP2GM123641, and R01GM107132. P.M.F. acknowledges support from the Beckman Foundation, the Sloan Research Foundation, and a McCormick and Gabilan faculty fellowship.

P.M.F. is a Chan Zuckerberg Biohub Investigator. K.B. acknowledges support from a TL1 Stanford Clinical and Translational Science Award to Spectrum (NIH TL1 TR 001084) fellowship, a Chemical Biology Interface (CBI) fellowship from the Chem-H Institute (Stanford University) and a NSF GRFP fellowship. C.J. M. acknowledges support from the Canadian Institutes of Health Research (CIHR) postdoctoral fellowship. T.C.S. acknowledges support from a NSF GRFP fellowship. B.C. acknowledges Stanford CAMPARE for support for a summer research rotation at Stanford University.

